# Single nucleotide variant at the αvβ_3_ integrin associated to Andes virus infection susceptibility

**DOI:** 10.1101/502591

**Authors:** C Martínez-Valdebenito, J Angulo, N. Le Corre, C Marco, C. Vial, JF Miquel, J. Cerda, GJ. Mertz, PA. Vial, M. Lopez-Lastra, M. Ferrés

**Author notes:** Corresponding author (MF).

## Abstract

**Background:** ANDV, agent of hantavirus cardiopulmonary syndrome, enters through integrin cell protein. A change from leucine-to-proline at residues 33 in the PSI-domain (L33P) inhibits ANDV recognition. We assessed the association between this human-variant and ANDV infection.

**Results:** We defined susceptible genotype to “TT” (coding leucine) and protective “CC” (coding proline). TT was in 89.2% (66/74) of a first cohort of ANDV-cases and in 60% (63/105) of exposed close-household contacts who remained unifected (*p*<0.05). Protective genotype was absent in all 85 ANDV cases in both cohorts and was present at 11.4% in exposed close-household contacts who remained uninfected. Logistic regression modeling to become a case had an OR 6.2-12.6 (*p*<0.05) in presence of TT and ANDV well-known risk activities. Moreover, OR of 7.3 was obtained when TT condition was analyzed for two groups exposed to the same environmental risk.

**Conclusion:** Host genetic background has an important role in ANDV-infection susceptibility in the studied population.

**Author summary:** Hantavirus is a worldwide infection known to cause two different diseases: the hemorrhagic fever with renal syndrome (HFRS) and hantavirus cardiopulmonary syndrome (HCPS or HPS), diseases seen in the old world and new world hantaviruses, respectively. Andes orthohantavirus (ANDV), is the etiological agent of HCPS in Chile and South of Argentina, and is the only hantavirus recognized to be transmited person-to-person. It has been suggested that β3-integrin is a cellular receptor for ANDV entry into cells. Here we try to understand the role of one human genetic variant of β3-integrin in susceptibility to ANDV-infection. We compared, exposed infected and non infected subjects, and their genotype differences regarding a β3 integrin variant that change a Leucine to Proline at PSI domain of the receptor. Leucine and proline aminoacids at the PSI domain, turn the cell susceptible or resistant to ANDV-infection, respectively. We were able to show genotype differences in cases and close-household contact that suggest differences in ANDV-infection susceptibility. Our results propose that the TT genotype, that codes to leucine, is a risk factor to become infected with ANDV, and the CC genotype that codes for proline at PSI domain, is a protective factor. (193/200)

## INTRODUCTION

The genus *Orthohantavirus*, members of the *Hantaviridae* family, are the etiological agents of two zoonotic diseases, known as hemorrhagic fever with renal syndrome (HFRS) and hantavirus cardiopulmonary syndrome (HCPS) (1,2). Andes orthohantavirus (ANDV) is the sole etiological agent of HCPS in Chile and south of Argentina, and its main reservoir is the long-tailed pygmy rice rat (*Oligoryzomys longicaudatus*) (3,4). Through September 29, 2018, a total of 1141 cases of ANDV have been reported in Chile, with a lethality of 30 to 35% (5). Transmission of ANDV to humans occurs mainly by exposure to aerosolized feces, urine and saliva of infected rodents. However, ANDV person-to-person transmission has also been reported in Chile and Argentina (5–7).

After environmental or interpersonal virus exposure, the incubation period for ANDV infections has been estimated to be between 7 to 39 days with an average of 18 days (6,8), while in the 2012 Yosemite HCPS-outbreak due to Sin Nombre virus, the median incubation period was 30.5 days with a range of 20-49 days (9). Four stages characterize clinical presentation of HCPS namely the prodromal, cardiopulmonary stage, diuresis and convalescent phases. The prodromal phase is characterized by fever, headache and myalgia. The cardiopulmonary phase presents with tachypnea and dry cough secondary to pulmonary edema that can quickly progress to respiratory failure, cardiogenic shock and death during this stage (1). After several days, spontaneous diuresis occurs among survivors of the cardiopulmonary stage. The convalescent phase has been poorly characterized (8). The first symptoms of the cardiopulmonary phase can progress rapidly to a severe disease with the need of mechanical ventilation (MV), the use of vasoactive drugs and even the use of extracorporeal membrane oxygenation (ECMO). Strikingly, some patients exhibit a mild disease with only minimal or total absence of oxygen supplement requirement (10,11).

Although risk factors for environmental and person-to-person transmission are well characterized, host factors that determine susceptibility to infection and disease severity are incompletely understood. In humans, pathogenic hantavirus such as ANDV replicate primarily in vascular endothelial cells. As such, differences in virus-cell affinity or the ability to attach to a known receptor may explain why viral attachment and entry is successful. Endothelial cells infected with ANDV induce the production of vascular endothelial growth factor (VEGF) followed by the downregulation of VE-cadherine, which leads to an increase in the microvascular permeability (12–14). Several surface proteins have been identified as mediators of virus entry into cells, (15) and in vitro studies have identified integrin as one of the main cellular receptors used by hantaviruses (16–18). The interaction between the envelope glycoproteins of ANDV and β_3_ integrin is mediated through the plexin-semaphorin-integrin (PSI) domain of the inactive integrin conformation (18).

It is noteworthy that the single nucleotide variant (SNV rs5918) in human β_3_ integrin is a missense substitution (NP_000203.2:p.Leu59Pro) within the 33^rd^ aminoacid of the PSI domain (NP_000203.2:P.LEU59 Pro) and has been shown to block human β_3_ integrin-ANDV interaction (14). This intriguing observation prompted us to design a study looking for genetic association analysis to address whether a link could be established between ANDV infection in Chilean patients and genetic variation in αVβ_3_ integrin SNV rs5918. If so, we predicted that individuals with SNV rs5918 leading to the (NP_000203.2:P.Leu59Pro) amino acidic substitution within the PSI domain would be less susceptible to ANDV infection. To evaluate this possibility, samples from three groups of individuals were analyzed. The first group consisted of healthy individuals known to be representative of the Chilean population (19). The second group consisted of a case-closehousehold contact population of individuals exposed to ANDV. This second group was further stratified to confirmed ANDV-infected index cases and their close-household contacts who remained uninfected during prospective followup. The third group consisted of close-household contacts who developed HCPS during prospective follow up (6).

## MATERIAL AND METHODS

### Study Population

Three sets of subjects were evaluated

1. **Chilean population (group 1)**: The first group, the general population, included DNA samples from 477 nonrelated and ANDV uninfected individuals obtained from a well-characterized DNA library harvested from a population considered to be representative of the Chilean population (19).
2. **ANDV cases and close-household contacts who remained unifected (group 2)**: The second group, HCPS cases and close-household contacts were both exposed to ANDV. Briefly, cases were 74 ANDV-infected individuals, confirmed through positive specific immunoglobulin M serology and/or by positive ANDV RT-PCR (20,21). The 105 close-household contacts were exposed to index cases and in some cases to common environmental risk factors but remained uninfected during five weeks of follow up. These close-household contacts slept in the same bed or had close contact with an ANDV-infected case during 30 days before and 7 days after the onset of HCPS symptoms. Both the HCPS cases and close-household contacts were enrolled between 2008-2014. Demographic and epidemiological data were collected for cases and contacts through a previously validated questionnaire (6).
3. **Household contacts who developed HCPS during prospective follow up (group 3)**: The third group included 11 subjects enrolled between 2002-2005 as healthy household contacts of ANDV cases who subsequently seroconverted and became ill during five weeks of prospective follow up [6]. DNA was available from 11 of the 14 household contacts who acquired ANDV infection in that study.

### Ethical statement

Approval for the use of all samples, data and research protocol design was obtained from the Ethical Review Board of the Facultad de Medicina, Pontificia Universidad Católica de Chile (Code 12-292 and 16-092). The participant or their legal representative sign a written consent form at the time of enrollment, this consent form was previously approved by the Ethical Review Board.

### DNA Extraction and Genotyping

Genomic DNA was extracted from cryopreserved blood samples using the MagNApure compact system (Roche^®^), according to manufacturer instructions. Genotyping of the rs5918 SNV was performed using a predesigned SNV assay with hydrolysis probes (ThermoFisher Scientific^®^, cat. n<u>º</u> 4351379). The amplification reaction was conducted in a Stratagene Mx3000P thermal cycler (Agilent Technologies), and the assignment of alleles was attributed automatically by the MXPro QPCR software version 4.10 (Agilent Technologies), as is described elsewhere (22) and manually reviewed by two independent investigators. To verify the correct assignment of alleles, control samples of each genotype (homozygote for each allele and heterozygote) were sequenced. Genotyping controls were added to each run and all samples were run in duplicates.

Subjects with TT-genotype (homozygote for the major allele) were defined as “susceptible” to ANDV infection, whilst genotypes CT and CC (heterozygote and homozygote for the minor alleles, respectively) as “protective” condition in the logistic regression model.

### Statistical Analysis

For descriptive analysis of each variable and odds-ratio (OR) calculation (95% confidence of interval), we used the software SPSS version 21 (SPSS, Inc., Chicago IL). The frequency distribution for each variable was compared using Fisher’s exact test for contingency tables or χ^2^, depending of the categorization of each variables, and a χ^2^ test was used to verify any discrepancies of SNV rs5918 distribution from Hardy–Weinberg equilibrium. Significance was considered p<0.05

A logistic regression model was used to assess environmental or person-to-person risks factors for hantavirus infection either in the presence or the absence of the “susceptible” or “protective” genotypes. To run the different multivariable models, for the genotype variable we collapsed the “CC” genotype (codified to proline or protective genotype) category, with the “CT” genotype, to avoid a zero value in the “CC” box for infected patients and run the regression model.

We calculate OR with a univariated modeling (OR crude) and 3 different strategies for the multivariant modeling. Briefly, the first included all the registered variables (OR^1^), the second (OR^2^) only included the variables that were statistically significant in the univariate model (crude OR), and the third (OR^3^) included only those variables described in the literature as risk factors involved in ANDV infection (6,10).

Additionally, we selected cases and household contacts that shared the same environmental exposure and evaluated the risk of ANDV infection for each genotype. To compare the severity of ANDV-induced diease and SNV genotype, we assign severe and mild categories according to the patient´s clinical outcome. A mild disease was characterized as febrile illness with nonspecific symptoms (eg. headache, myalgias, chills, gastrointestinal symptoms) with no or minimal respiratory compromise. Severe cases were characterized for a rapid and progressive imparied lung function with mechanic ventilation and vasoactive drugs. Severe and mild were compared by χ^2^ test, using Graphad Prism version 7.04.

## RESULTS

### Genotypes distribution in general population

The genomic DNA from 477 healthy individuals obtained from a well-characterized DNA library considered to be representative of the Chilean population (19) were analyzed. The frequency for the rs5918 TT, TC and CC genotypes were 84.5%, 13.4%, and 2.1%, respectively. The SNVs rs5918 genotype showed to be in a Hardy Weinberg equilibrium (χ^2^ tests *p*≥ 0.1) (Fig. 1).

**Figure 1.**
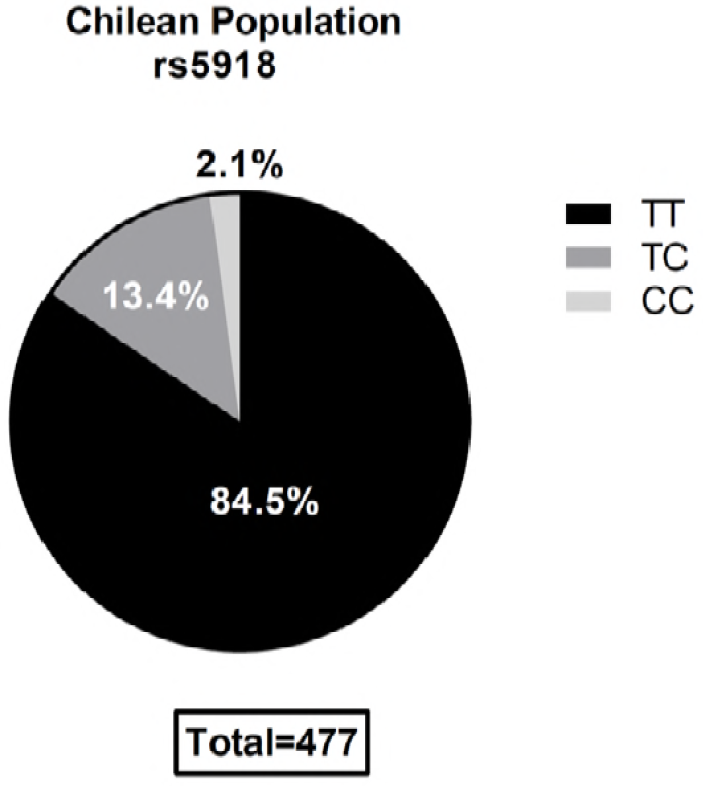
SNV rs5918 genotypes distribution within the Chilean population. The TT genotype is the homozygote that codes to leucine aminoacid at 33 position of the PSI integrin domain. The genotype CC is the homozygote that codes to proline amino acid at same position, reducing dramatically the ANDV recognition of in *ex vivo* models (14). SNV were in Hardy-Weinberg equilibrium (*p*>0.05).

### Analysis of SNV rs5918 distribution among study Group 2 (cases and close-household contact and study Group 3 (11 close-household contacts infected)

The TT genotype had a higher distribution among ANDV-infected subjects (89.2 %) than among the close-household contacts (60%) (Fig.2). The protective CC genotype was absent from all ANDV-infected cases, but present (11.4%) in the exposed but not infected close-household contacts (*p*<0.05). The TC genotype was found only in 10.8% of the ANDV-infected cases but in 28.6% of the close-household contacts who remained uninfected (Table 1). Among the 11 household contacts who acquired ANDV infection, five had the TT genotype, while 6 had the CT genotype and none had the CC protective genotype.

**Figure 2.**
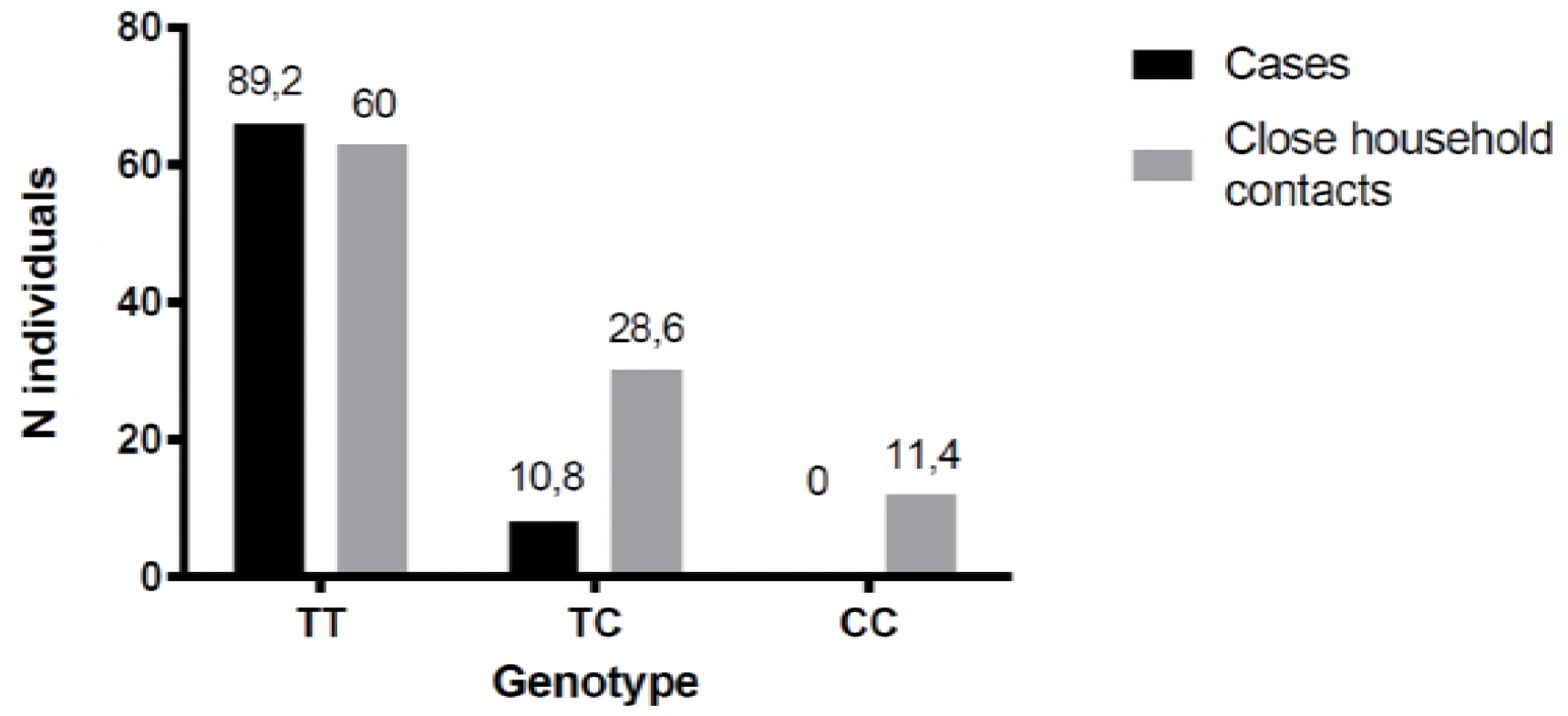
SNV rs5918 genotypes distribution in Cases and close-household contact. The cases and household contacts were grouped according to SNV 5918 genotype. The total number of each population was defined as 100% and the percentage of individuals according to each genotype is indicated. (*p*>0.05).

**Table 1.**
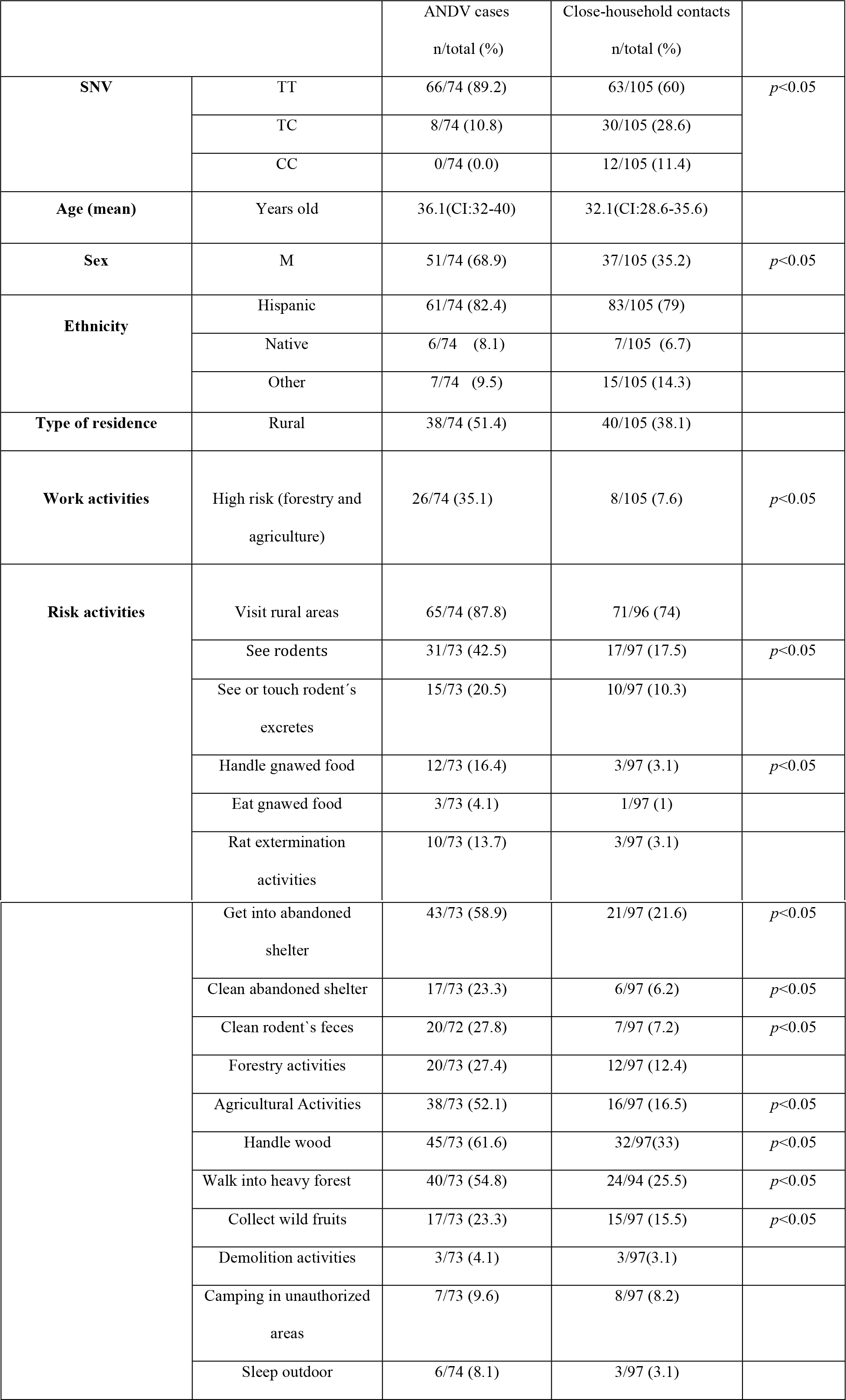
SNV rs5918 genotypes and risk variables distribution in ANDV infected cases and uninfected close-household contacts

Variables previously documented as risk factors for ANDV-infection such as cleaning or entering into abandoned places, wood handling, farm and forestry activities, or living in rural area, among others, showed clear differences between the ANDV-infected cases and close-household contacts that remained uninfected. (Table 1).

### ANDV infection risk assessment and risk models for SNV rs5918 genotype and infection in cases and close-household contacts

The risk of ANDV-infection was assessed based on the presence of the TT (susceptible) versus the CC/CT genotypes (defined as protective for the model) and the environmental variables. The crude OR for the existence of TT genotype and ANDV-infection was 6.2 (CI:2.7-14.1) (*p*<0.05). When demographic and all the exposure variables were added to the multivariable logistic model, the OR^1^ for TT genotype increased to 19.7 (CI: 3-131). Finally, when we only included variables with a significant crude OR (model 2) and variables that are well recognized in the literature as relevant for ANDV-infection (model 3), we obtained an OR^2^ and OR^3^ for the TT genotype of 12.6 (CI 2.9-55.3) for both (Table 2) *p*<0.05.

**Table 2.** OR for three logistic regression models linking genotypes and risk factors for ANDV acquisition. Data and numbers were shown at table 1.

|  | OR crude (CI) |  | OR<sup>1</sup> |  | OR<sup>2</sup> |  | OR<sup>3</sup> |  |
| --- | --- | --- | --- | --- | --- | --- | --- | --- |
| **SNV (TT)** |  | 6.2 (2.7-14.1)* |  | 19.7 (3-131)* |  | 12.6 (2.9-55.3)* |  | 12.6 (2.9-54.5)* |
| **Sex (M)** |  | 4.1 (2.2-7.7)* |  | 1.3 (0.3-5.1) |  | 1.5 (0.4-4.7) |  | 1.3 (0.4-4.4) |
| **Type of residence (R)** |  | 1.7 (0.9-3.1) |  | 0.1 (0.02-0.8) |  | ND |  | ND |
| **Work activities** |  | 8.8 (3.4-22.5)* |  | 21 (2.5-178)* |  | 7.9 (1.6-39.4) |  | 8.1 (1.6-40.6) |
| **Visit rural areas** |  | 2.5 (1.1-5.9)* |  | 0.4 (0.05-3.6) |  | 1.4 (0.4-5.1) |  | 1 (0.2-5.6) |
| **See rodents** |  | 3.5 (1.7-7)* |  | 1 (0.2-5.4) |  | ND |  | ND |
| **See or touch rodent's excretes** |  | 2.2 (0.9-5.4) |  | 0 |  | ND |  | ND |
| **Handle gnawed food** |  | 6.2 (1.7-22.7)* |  | 8.3 (0.3-248) |  | 1 (0-10.5) |  | ND |
| **Eat gnawed food** |  | 4.1 (0.4-40.4) |  | 1.9 (0.04-87) |  | ND |  | ND |
| **Rat extermination activities** |  | 5 (1.3-18.8)* |  | 4.4 (0.2-88) |  | 1.2 (0-19.2) |  |  |
| **Get into abandoned shelter** |  | 5.2 (2.6-10.1)* |  | 11 (2-60)* |  | 3.3 (1-10.7) |  | 4 (1.2-13) |
| **Clean abandoned shelter** |  | 4.6 (1.7-12.4)* |  | 0.6 (0.1-5.4) |  | 0.8 (0.1-5.4) |  | 0.8 (0.1-4.6) |
| **Clean rodent's feces** |  | 5 (2-12.5)* |  | 4.5 (0.3-68) |  | 1.7 (0.2-17.1) |  | 2.4 (0.4-14.7) |
| **Forestry activities** |  | 2.7 (1.2-5.9)* |  | 0.1 (0-0.6) |  | 0.2 (0-1) |  | 0.2 (0-1.2) |
| **Agricultural Activities** |  | 3.1 (1.5-6.4)* |  | 0.2 (0-1.5) |  | 0.9 (0.2-3.3) |  | 0.8 (0.2-3.1) |
| **Handle wood** |  | 3.2 (1.7-6.2)* |  | 3.4 (0.7-16.3) |  | 2.8 (0.8-9.8) |  | 2.5 (0.7-9.4) |
| **Walk into heavy forest** |  | 3.5 (1.8-6.8)* |  | 38 (4-349)* |  | 6.6 (1.8-23.4) |  | 7.2 (2-25.3) |
| **Collect wild fruits** |  | 1.7 (0.8-3.6) |  | 3.7 (0.6-22) |  | ND |  | 1.8 (0.5-6.8) |
| **Demolition activities** |  | 1.3 (0.3-6.8) |  | 0.3 (0-5) |  | ND |  | 0.5 (0-8.5) |
| **Camping in unauthorized areas** |  | 1.1 (0.4-3.4) |  | 3.2 (0.3-33) |  | ND |  | 1.2 (0.2-8.7) |
| **Sleep outdoor** |  | 2.8 (0.7-11.4) |  | 0.5 (0-6) |  | ND |  | ND |

\* $p < 0.05$ ; CI: Confidence interval 95% was used. <sup>1</sup> All variables were included; <sup>2</sup> Only statistically significant variables were included; <sup>3</sup> Variables selected according to literature. \*ND= not done

### ANDV infection risk assessment among cases and uninfected close-household contacts exposed to the same risk activity

In order to rule out the difference in exposure of cases and close-household contact,we selected the two more frequent risk activities shared between ANDV-infected cases and close-household contacts who did not became infected and assessed the OR of carrying the susceptible and protective genotypes of SNV rs5918. When we linked accessing an abandoned building, the susceptible genotype TT was present in 90.7 % (39/43) of cases and in 57.1 % (12/21) of the close-household contacts. Otherwise, for wood handling the TT trait was present in 84.4% (38/45) of cases and 59.4% (19/32) of close-household contacts. The OR for ANDV infection in presence of TT genotype for each activity was 7.3 (1.9-28) and 3.7 (1.3-10.8), respectively (Table 3) *p*<0.05

**Table 3.** Distribution of SNV rs5918 genotypes in ANDV cases and uninfected close-household contacts exposed to the same risk activity.

| Genotype | Access to abandoned places<sup>1</sup> |  | Handle wood<sup>2</sup> |  |
| --- | --- | --- | --- | --- |
|  | Cases n (%) | Close-household contacts n (%) | Cases n (%) | Close-household contacts n (%) |
| Susceptible (TT) | 39 (90.7) | 12 (57.1) | 38 (84.4) | 19 (59.4) |
| Protector (TC/CC) | 4 (9.3) | 9 (42.9) | 7 (15.6) | 13 (40.6) |
| **Total** | **43** | **21** | **45** | **32** |

<sup>1</sup> OR: 7.3 (IC: 1.9-28); <sup>2</sup> OR: 3.7 (IC: 1.3-10.8)

The protective genotype (CC or CT) associated to each activity was present in 9.3% (4/43) and 15.6% (7/45) in ANDV-infected cases and in 42.9% (9/21) and 40.6% (13/32) for uninfected close-household contacts.

### Severity of ANDV-induced disease and SNV rs5918 genotype distribution

We classified the cases in mild or severe disease according to the patient´ final clinical outcome. Fig. 3 show no differences in the genotype distribution between severe and mild disease with a p>0.99.

**Figure 3.**
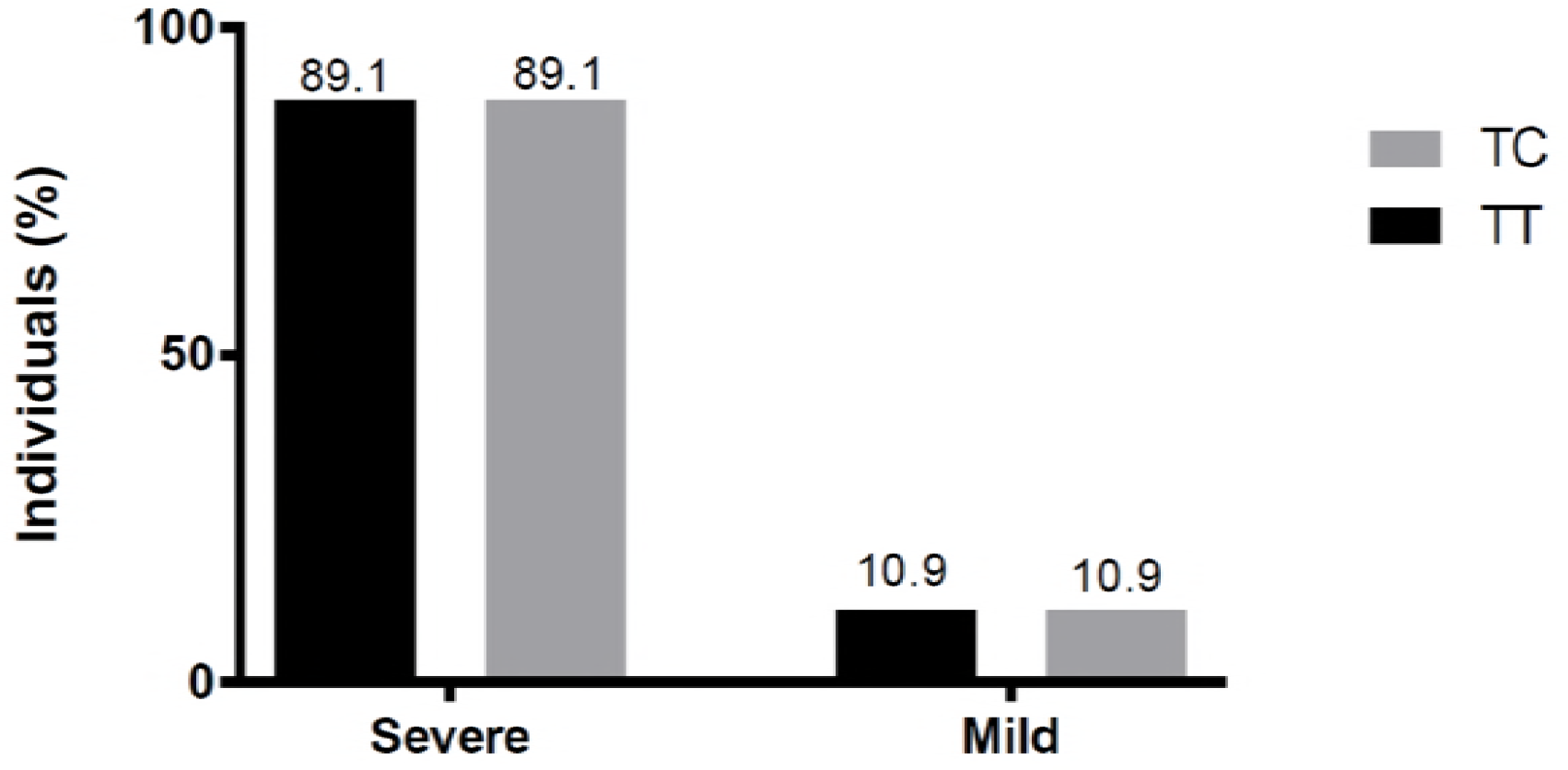
Genotype distribution in subjects with severe or mild disease. Severe patients were 4 TT and 4 CT, meanwhile for mild disease TT and CT were present in 33 subject.

## DISCUSSION

Recent studies have linked the severity of ANDV-illness to genetic factors^14^. Herein we have sought to address whether the risk of infection could also be associated with host variants. In that direction, a link between SNVs rs5918 and ANDV-infection has been suggested by *in vitro* studies (14,18). In the current study we found that SNV rs5918 have an important role in the infectious process of ANDV. The ANDV-infected cases (74) showed a frequency of 89.2% regarding the susceptible genotype of the SNVs rs5918 while the the protective CC genotype was totally absent. Furthermore, none of the 11 household contacts who acquired ANDV infection during prospective follow up had the CC genotype. In addition, the protective CC genotype was present in 11.4% of the exposed household contacts that did not become infected by ANDV. All these finding support the conclusion that the rs5918 TT genotype confers susceptibility to ANDV infection while the rs5918 CC genotype is protective.

Nevertheless, is important to mention the remarkable difference in frequency of CC genotype in the household contact compared to the Chilean population and others reports regarding to rs5918, were the frequency for this genotype is not more than 2% (23). As we mention the close-household contacts population were exposed to environmental risk factors of ANDV-infections and particularly to person-person transmission. These population were the sexual partner of the case, or parents or children of the cases and therefore blood related for this last two situations. This can explain the differences founded in this particular cohort compared to Chilean population and previous reports of rs5918 (22).

Several authors have proposed that integrin αvβ_3_ is the main receptor of entry for pathogenic hantaviruses such as ANDV and suggested its role in the pathogenesis of the disease (16,17). *Lui et. al. 2009* showed that a variant in the HPA-3b allele (I843S) (integrin α_IIb_β_3_) resulted in more severe clinical HFRS (24). In addition, NP_000203.2:P.Leu59Pro substitution, in PSI domain of β_3_, is capable of directing autoimmune responses to β_3_ integrins from blood containing a different HPA type. This response to β_3_ results in two autoimmune diseases, that display vascular permeability and acute thrombocytopenia similar to hantavirus pathogenesis (25). Thus, it was plausible to suspect that SNV rs5918 in β_3_ could also be associated with the ability of ANDV to infect or to cause disease upon infection.

In a recent report, the association between SNV rs5918 and hantavirus infection was studied in a Chinese population. In this study authors failed to establish an association between the SNV rs5918 and the susceptibility to infection with Hantaan and Seoul viruses, both responsible for HRFS (26)

Notwithstanding the above, in this study we decided to evaluate the association between SNV rs5918 and ANDV-infection due to differences between old world and new world hantaviruses (1,12), in the illnesses in humans, and genetic differences between the Chinese and Chilean populations. The Chilean population, which includes ancestral contributions from Europe, Native Americans, and a minor African component (27), differs sharply from what is expected from the Asian cohort. In contrast to what was shown for a Chinese population, results presented herein suggest that SNV rs5918 links with the ability of ANDV to infect Chilean subjects. This may suggest that the ethnic background should be accounted for when establishing genetic studies. It should be noted however that genotype TT is prevalent among general Chilean population, an observation that could bias our conclusions. However, in support of our conclusion is the observation that the protective CC genotype was absent in the ANDV-infected cases and more prevalent in the close-household contacts.

*In vitro* assays, establish the critical role of leucine at site 33 of the β integrin PSI domain for ANDV infection (14). SNV rs5918 in the *ITGB_3_* gene defines the Leu33Pro substitution. Genotype TT results in a leucine at position 33 that is expected to facilitate the entry of the virus (12,14), while the CC genotype implies a proline, that hinders binding of the ANDV-glycoproteins to the integrin receptor (14). Based on these observations in the present study, we evaluated the potential association between the risk of an ANDV-infection and the SNV rs5918 in the aim of understanding how the hosts genetic background impacts on the distribution of an infectious disease.

As we mention, the environmental risk factors for ANDV acquisition include living in rural areas, working in forestry or agriculture, and recreational activities such as camping activities or sleeping outdoors. Through logistic regression models we weighted the SNV rs5918 impact in presence of a hantavirus risk activities assessing the infection as a final outcome. The OR for susceptible TT genotype was statistically significant in all applied models, highlighting the relevance of this β_3_ integrin variant for ANDV-infection. For the modeling purpose, we used the CT and CC genotype condition as protective. Nonetheless, data for individuals exhibiting SNV rs5918 heterozygosity are complex to interpret and should be regarded with caution. The rs5918 CT genotype is present among both ANDV-infected cases, close-household contacts and the general population in similar proportion. For the 11 prospective cases analyzed it would be expected that the TT were the most prevalent genotype, however the CT distribution was higher, maybe due a the small subject analyzed. However, what is still unclear is the phenotype expression, at position 33 of the β_3_ protein, in the individuals that are heterozygous for the rs5918. Thus, without the direct amino acid sequence that is expressed in this cases, we cannot exactly determine if the CT genotype represents a susceptible or protective condition. While, SNV rs5918 showed a high OR in the 3 different regression models emphasizing the role in the ANDV-infection in humans, it would also be interesting to determine if this interaction between virus-cell is the only characteristic responsible for the ability of ANDV to cause disease in Syrian hamster animal model or the only responsible characteristic for ANDV to be transmitted person-to-person (14). Recently, Jangra et al. 2018 show the role of PCDH1, cadherine superfamily, as a critical host factor for cell entry and infection in a Syrian hamster model(30).

One general characteristic of human infection is that not all individuals exposed to a pathogen became ill. To address this, different genetic markers have been associated to this broad range of susceptibility to infectious diseases. A good example is tuberculosis in African population of Gana, Gambia and Malawi, where a SNV (rs4334126) show a OR of 1.19 to develop tuberculosis illness. This SNV is located in a conservative region of the 18q11.2 chromosome, suggesting a possible regulatory effect on an unknown gene (28,29). Here we showed that cases and uninfected close-household contacts exposed to the same risk activities have statistical difference in frequency distribution for NP_000203.2:P.Leu59Pro variant in rs5918 SNV with OR of 7.3 and 3.7 for becoming infected, suggesting a clear difference in the susceptibility to ANDV infection.

In summary, either with the tested logistic regression models and the SNV rs5918 distribution in population at same risk activities, we were able to show the role of SNV rs5918 in susceptibility to ANDV-infections. Nevertheless, other receptor and factor contributed to the global host susceptibility.

Our work highlights the relevance of the genetic background of the host in the susceptibility to an infection and help to understand why two equally exposed individuals have different outcomes of infection.

## ACKNOWLEDGMENTS

In addition to the authors we thanks to members of the Hantavirus Study Group in Chile who contributed to patient enrollment and follow-up, sample collection and analysis, and data management. We also, thanks to Francisca Valdivieso MD for her work in the development of the protocol.

